# Assembly of Yeast Spindle Pole Body and its Components Revealed by Electron Cryo-Tomography

**DOI:** 10.1101/442574

**Authors:** Sam Li, Jose-Jesus Fernandez

**Author notes:** Correspondence: Sam Li MC 2240 University of California, San Francisco San Francisco, CA 94158, USA.

## Abstract

Spindle pole body (SPB) is the microtubule organizing center (MTOC) in yeast. It plays essential roles during many cellular processes, ranging from mitosis to karyogamy. Here, we used electron cryo-tomography (cryo-ET) and image processing to study SPB and its component purified from the budding yeast. The 3D images and models of SPB at various cell cycle stages were reconstructed by cryo-ET and were analyzed. The results reveal SPB as a cylindrical shaped structure composed of multiple layers. The central layers are arranged with a degree of crystalline order. By using subtomogram averaging methods, we studied the purified “sheet” from over-expressed Spc42p. Our analysis of the SPBs and its components provides new insights into the assembly of this organelle and its cellular function as an MTOC.

## Introduction

The spindle pole body (SPB) is the sole microtubule organizing center (MTOC) in budding yeast, and is functionally equivalent to the centrosome in higher eukaryotes. The SPB is a cylindrical shaped organelle, permanently embedded in the nuclear envelope. The nuclear and cytoplasmic microtubules (nMTs and cMTs) are attached to either end of the SPB. It plays critical roles in a number of cellular events, including chromosome segregation during mitosis and meiosis, nuclear congression in karyogamy, spindle positioning and nucleus migration. It is also essential for forming prospore membrane in meiosis [1]. In addition, the SPB transiently recruits cell cycle regulators such as the mitotic exit network (MEN) components to the site to ensure the integrity of cell division [2, 3]. The molecular composition of the SPB has been identified and characterized by a combination of biochemical and genetic approaches. Together, these studies have provided a comprehensive list of major components of the SPB and their roles in the SPB assembly and functions (for reviews, see [4, 5]). In parallel, a number of approaches have been applied to localize the SPB components and to elucidate the molecular organization of the entire organelle. These include immuno-EM, electron tomography [6, 7], fluorescence resonance energy transfer (FRET) experiments [8] and integral bioinformatics approach [9]. Furthermore, to date a number of high resolution structures of individual component and complexes of SPB have been solved by X-ray crystallography [9–12] and electron cryo-microscopy (cryo-EM) [13].

However, our understanding of the SPB assembly is far from complete. Here we have used cryo-electron tomography (cryo-ET) on enriched SPBs from budding yeast *S.uvarum,* similarly to an earlier study [7], but improved technology and methods make increased resolution possible.

## Results and Discussions

### The SPB is Composed of Multiple Layers

We first studied the overall structure of this organelle by cryo-ET. Previous cryo-EM studies on enriched SPBs used 2D projections [7], here we have used 3D cryo-ET to improve the resolution. A raw image is shown in Fig. 1A and a slice from its reconstructed 3D volume is in Fig. 1B. The entire tomographic tilt series and its 3D reconstruction can be seen in supplemental videos 1 and 2. Compared to previous EM studies, the improved resolution by cryo-ET is evident. At least 8 different layers can be seen (Fig. 1C). The central core layers, IL2, CP1 and CP1, are particularly well-ordered as shown from the striations visible in the edge-on view of these three layers (Fig. 1, B and C). The IL2 layer will be discussed in more detail later. This layer has previously been shown to have hexagonal packing in heparin-extracted SPBs [7], and is ordered in intact cells [6]. Interestingly, the minus ends of some nMTs are closer to the CP2 layer than others (Fig. 1B), this might be due to flexibility of the Spc110p coiled coil, which would help accommodate a bundle of nMTs within a limited surface area and also allow nMTs to grow at various angles relative to the SPB’s longitudinal axis thus interdigitating in paired SPBs and thereby facilitating spindle formation.

**Figure 1.**
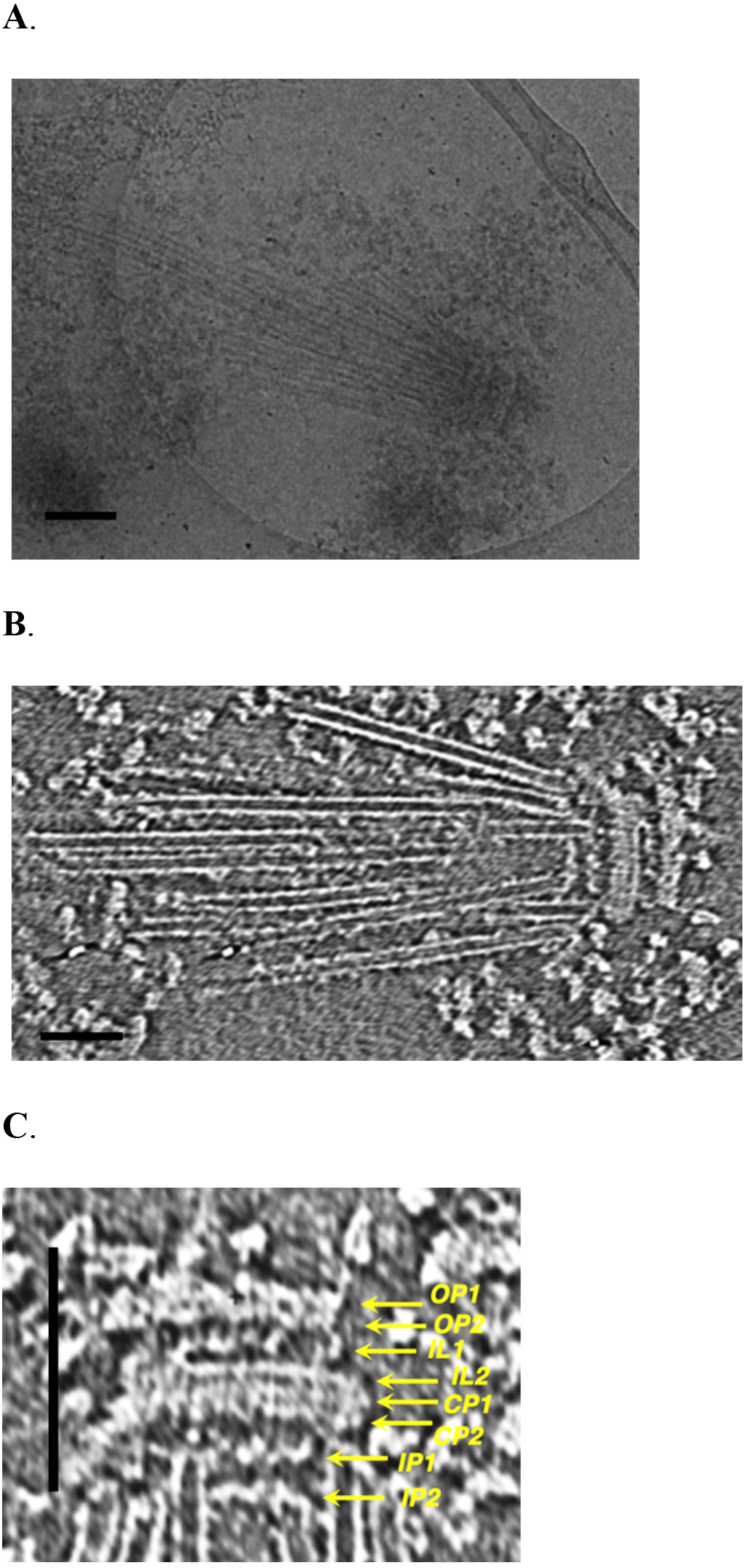
Electron cryo-tomography of the purified yeast spindle pole body. (A) A low-dose cryo-EM image of a SPB embedded in vitrified ice. (Scale, 200 nm)
(B) A slice from a 3D reconstructed tomogram shows a SPB along with bundle of nuclear microtubules. (Scale, 100 nm)
(C) The SPB is composed of eight layers. The image is rotated from the original so that the orientation of the SPB is vertical from the cytoplasm (top) to the nucleoplasm (bottom). A vertical scale indicates 200 nm. The measured spacing between neighboring two layers are as following: OP1-OP2, 17.6 nm; OP2-IL1, 27.4 nm; IL1-IL2; 21.6 nm; IL2-CP1, 19.6 nm; CP1-CP2, 15.7 nm; CP2-IP1, 35.3 nm; IP1-IP2, 37.2 nm. A histogram depicting eight layers and their peak distances is shown in Supplementary Fig. S1.

We can identify the probable positions of SPB components within these layers, summarized in (Fig. 6). These are based on previous immunoEM results, analysis of deletions and of purified SPB cores [14–19], also two-hybrid, genetic, together with complex isolation and coimmunoprecipitation results [20–23].

### Spindles in Early Mitosis

The SPBs were purified from asynchronous cell. This allowed us to capture SPBs in different cell cycle stages, including entire bipolar spindles at different time points in mitosis. Here, we present two examples of mitotic spindle. Based on the spindle length and morphology, they are likely to be in early metaphase [24, 25].

The first mitotic spindle has a pole-to-pole distance of ~1.0 micron (Fig. 2A, Supplemental video 3). The top-right SPB has 32 nMT attached, while the bottom-left SPB has 25 nMT attached. The length of MT varies substantially from 63 nm to 745 nm. We found 5 pairs of antiparallel MTs that are close enough to make potential crosslinking interactions (within 35 nm, highlighted in red in the model). Within them, a long overlap stretches ~180 nm in the middle of spindle. Another antiparallel pair makes a short overlap about 22 nm long. The remaining three pairs are MTs that mutually crossing at an angle, making only point contacts. The overall overlap length is 291 nm.

**Figure 2.**
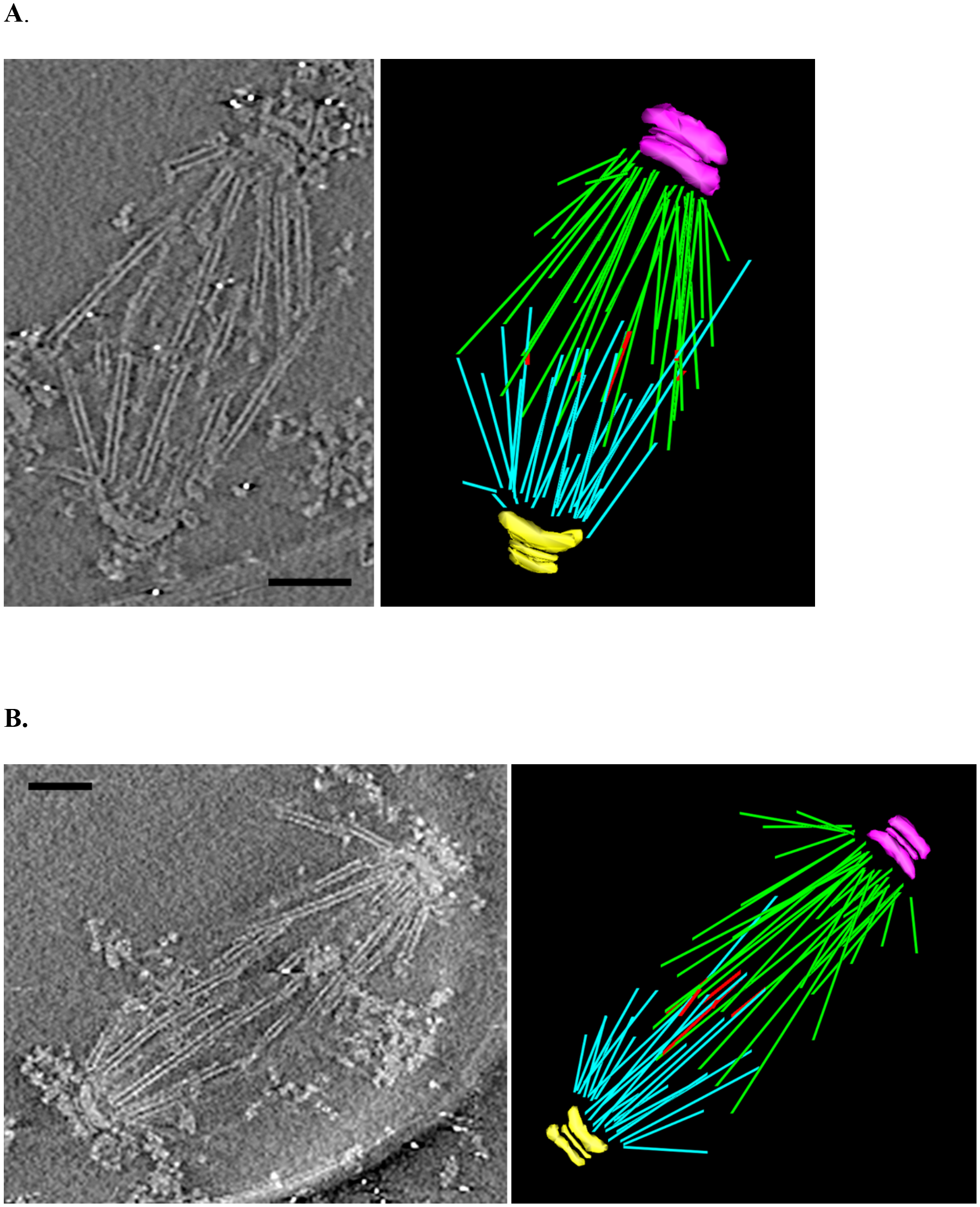
Two reconstructed mitotic spindles and their 3D models. (A) A 3D reconstruction of mitotic spindle in metaphase. The panel on the left is a central slice from a reconstructed 3D volume. One the right is a 3D model based on the reconstruction. In the model, the two SPBs are colored in pink and yellow and their associated nMTs are colored in green and cyan, respectively. The overlapped regions between antiparallel MTs are highlighted in red. (Scale, 200 nm)
(B)A second mitotic spindle in metaphase. As in (A), the left panel is a central slice from the reconstructed 3D volume. The right panel is a 3D model built based on the reconstruction. In the model, two SPBs are colored in pink and in yellow respectively and their associated nMTs are in green and cyan, respectively. The overlapped regions between antiparallel MTs are highlighted in red. (Scale, 200 nm)
(C) An example of crosslink between antiparallel MTs in the second mitotic spindle.
Inset: a yellow arrow indicates a crosslink between antiparallel MTs. The length of the crosslink is ~33 nm. (Scale, 200 nm)
(D) Histograms show the unimodal distribution of microtubule length in the above two mitotic spindles.

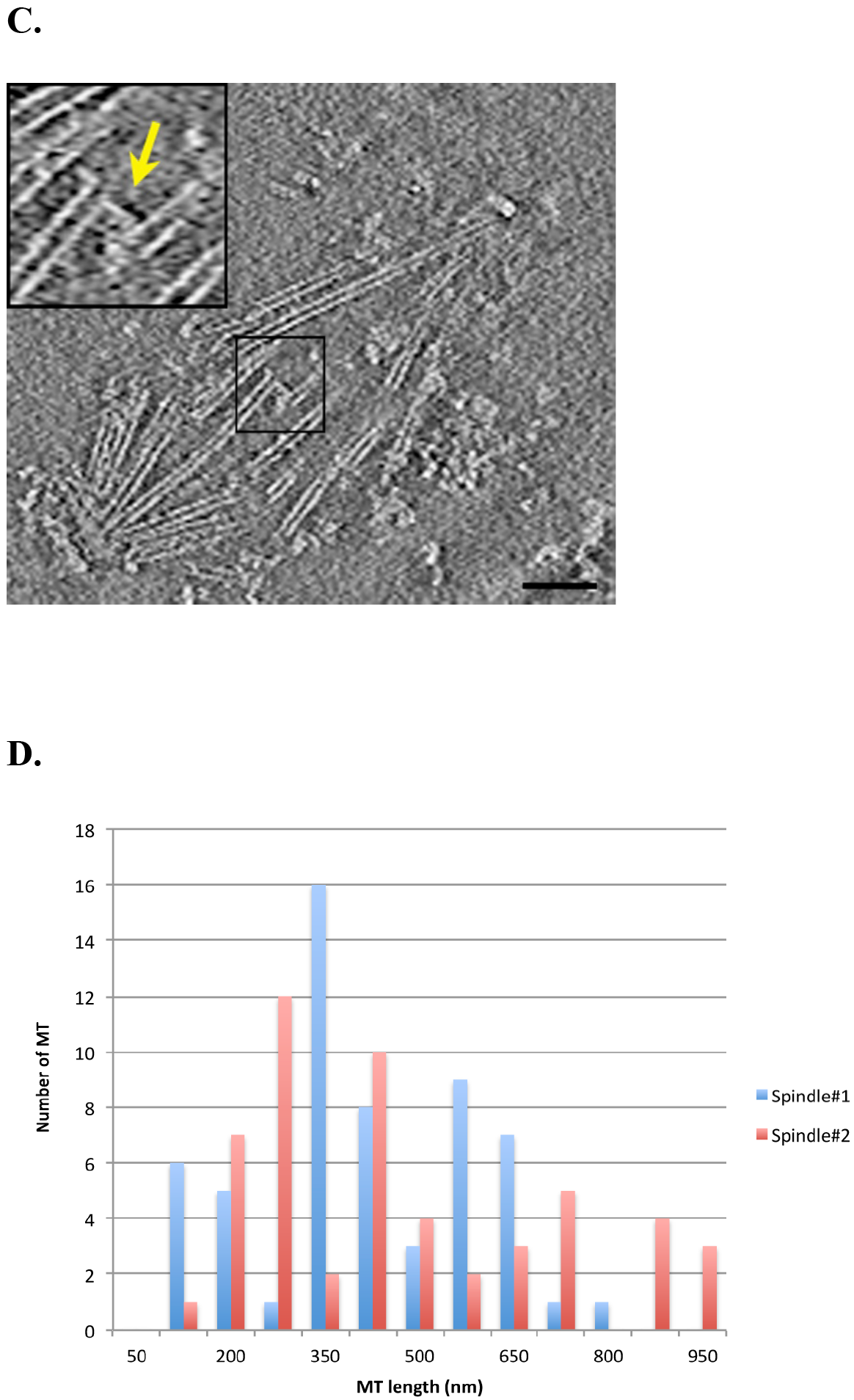

In a second reconstructed spindle (Fig. 2B, Supplemental video 4), the pole-to-pole distance is 1.5 microns, indicating that mitosis in this cell has proceeded further compared to the first one. The top-right SPB has 29 nMTs attached and the bottom-left SPB has 24 nMTs attached. The length of nMTs ranges from 100 nm to 937 nm. We found a total of 7 pairs of antiparallel MTs that are within 35 nm distance. All are clustered near the spindle center with different overlap length, ranging from 32 nm to 346 nm (highlighted in red in the model). The overall overlap length is 879 nm, more substantial than the first one. An example of a cross-bridge between a pair of antiparallel MTs is shown in Fig. 2C.

Since the spindles were embedded in ice but without cryosectioning, potentially they could be slightly compressed in a direction perpendicular to the direction of ice layer during the vitrification process. This might have brought MTs to vicinity that appears as overlapping MTs. However, the fact that all overlaps are clustered in the center of the spindle, and the antiparallel overlapping MTs are only a small percentage of total number of MTs in the spindle, makes us to believe that majority of the overlapping MTs that we observe are genuine structures, especially those near-perfect antiparallel pairs, where crosslinking structures have been observed. Even though the contacts are moderate compared to the interpolar MTs in anaphase, the crosslink is strikingly strong enough to endure multiple purification steps and preserve the overall structure of bipolar spindle. The extent of overlaps is also consistent with previous observations *in vivo* [6, 25, 26]. These crosslinks, presumably by Ase1p, a yeast homolog of human protein PRC1, stabilize anti-parallel microtubules in a MT length dependent manner and ensure the integrity of the spindle [27, 28]. As mitosis proceeds further, these crosslinked MTs will potentially form interdigitating interpolar MTs that establish the midzone at onset of anaphase and maintain spindle stability and orientation over long distances [29].

To further analyze the overall length distribution of MT in these two spindles, we plotted the histograms of MT with respect to their lengths (Fig. 2D). Unlike the bimodal distribution of MT in anaphase, here both histograms show a unimodal distribution of the MT length that is peaked at ~350 nm, indicating there is no shortening of the kinetochore-MTs or elongation of the interpolar MTs at these stages. This is consistent with previously reported MT length distribution in metaphase spindles in yeast [6, 25, 26, 30].

### The SPB Bridge

The SPB is duplicated once per cell cycle similar to the centrosome in higher eukaryotes. The bridge, which is attached to one side of the SPB, has a crucial role in SPB duplication. It consists of a thin rectangular cytoplasmic layer, called the outer layer, which probably contains the centrin-binding protein Sfi1p [11, 31]. This outer layer is closely associated with darkly stained inner and outer nuclear membrane, where the transmembrane proteins Kar1p [32] and Mps3p [33] are localized. In single SPBs the bridge is shorter and called the half-bridge [34]. When duplication starts the bridge doubles in length [11, 34], then Spc42p, Spc29p, Cnm67p and Nud1p assemble at the distal end of the elongated bridge [19], to form first a satellite then eventually a new SPB. We were interested in using cryo-ET to establish more precisely where the bridge outer layer, and presumably Sfi1p, was connected to the SPB. Thus paired SPBs, which would have just undergone duplication and were still connected by a bridge, were examined. Fig. 3A and Supplemental video 5 show a reconstruction of paired SPBs and its model. A bridge structure connects these two SPBs over a distance of 110 nm, consistent with previous reported bridge lengths [11]. To look at the bridge-SPB connection in more detail, we took the 9 reconstructed paired SPBs, out of a total of 150 SPB reconstructions, and for each reconstructed volume, we selected the central slices showing the most prominent bridge structure, then cut this in half and rotated one SPB through 180° to superimpose it on its pair and subjected these paired SPBs and their attached bridge structures to 2D alignment and averaging. The averaged result, as shown in Fig. 3B, resolved multiple layers in the SPB, and showed a clear bridge outer layer connected to CP1. Note that these enriched SPBs are detergent-extracted so the nuclear membrane part of the bridge is missing. The bridge outer layer is slanted at about 10° relative to the CP1 layer, this is presumably to accommodate the curvature of the nuclear membrane against the side of the SPB. Bbp1p is localized to this linkage site where it interacts with the core SPB component Spc29p and the transmembrane components Mps2p [35] and Mps3p [36]. Ndc1p and Nbp1p also participate in these interactions [37].

**Figure 3.**
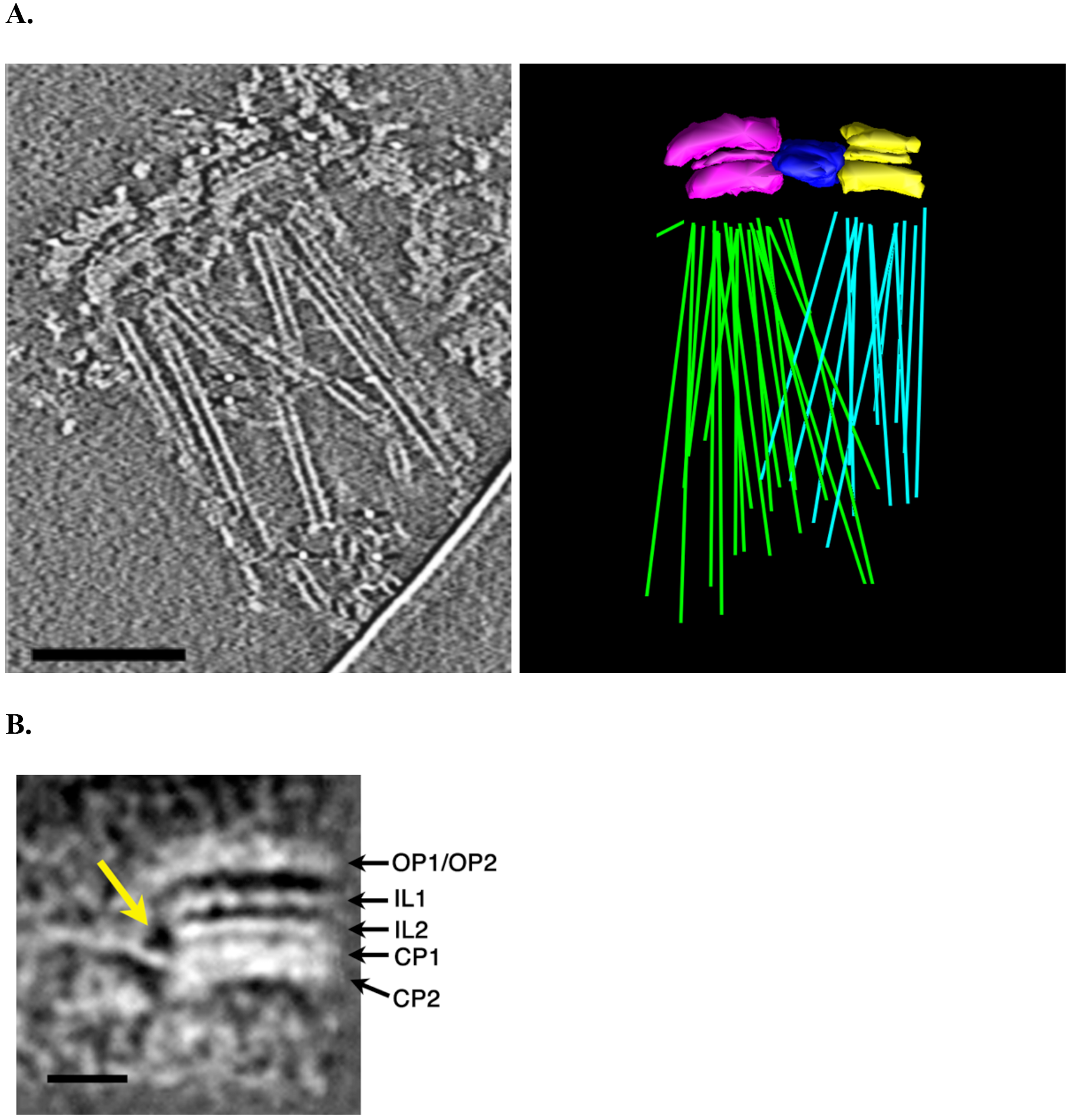
A pair of SPBs in duplication. (A) A central slice from a 3D reconstruction and a model showing a pair of SPBs in duplication. One SPB is larger and is colored in pink and the other smaller SPB is colored in yellow. Two bundles of nMTs are colored in green (22 nMTs) and cyan (13 nMTs) respectively. The bridge structure that links the two SPBs is colored in dark blue. (Scale, 200 nm)
(B) The 2D averaged duplicating SPB shows the bridge links to the CP1 layer of the SPB. Total 18 SPBs from 9 duplication pairs were used in the average. The yellow arrow indicates the linkage between the SPB and the bridge. (Scale, 50 nm)

In all 9 reconstructed pairs of SPB studied above, all eight layers can be discerned vertically in the SPB, suggesting Spc72p and Spc110p have already been recruited to the new SPB at this point of the cell cycle. Interestingly, we observed lateral size difference between two SPBs within each pair. Based on the diameter of the IL2 layer, one SPB, presumably the mother SPB, has larger diameter (152 ± 9 nm, mean/standard deviation) than the other SPB (121 ± 17 nm, mean/standard deviation) (the pairwise student’s t-test *p* value = 0.00003). The measured sizes are consistent with previous report in diploid cell where two major lateral-size classes have been found, even though in this case the measurement was not restricted to the duplicating SPB pair [7].

### Assembly of Sheets by Over-expression of Spc42p

When Spc42p, one of the core components of the SPB, is overexpressed in yeast it forms an ordered 2D sheet [38]. This grows out from the side of the SPB where Spc42p is localized [15], and associates with nuclear membrane on one side and endoplasmic reticulum on the other. Since these two membranes are equivalent, this suggests the Spc42p sheets are symmetric, that is composed of two layers of Spc42p exposing identical surfaces on either side. The sheets can be purified and consist of Spc42p only [38]. A more detailed structural examination of the Spc42p sheets showed them to be crystalline with a hexagonal pattern, a similar pattern was found in the central IL2 layer in heparin extracted SPB cores where Spc42p is localized, demonstrating that both assemblies share the same architecture [7]. In order to gain further insight into the assembly of Spc42p, we studied these purified sheets by cryo-ET and subtomogram averaging. Fig. 4A shows a cryo-EM image of a sheet and a central slice from a cryo-ET reconstructed 3D volume. Both clearly show ordered assemblies. A Fourier transform of the reconstructed central slice exhibited a characteristic hexagonal pattern (Fig. 4B). The first order reflection is at a spacing of 132 Å, which is close to the previously reported value of 126 Å measured from positively stained specimens [7]. However, in the reconstructed tomogram the reflections in the Fourier transform are limited only to the first order and are smeared, indicating local disorder of the crystallinity. We attempted to correct this distortion by subtomogram alignment and averaging, where local small patches were treated as repeat units followed by correction for their shift and rotation individually. A total of 660 subtomograms were used for alignment. The resulting average represents the repeat unit for the sheet (Fig. 4C). Overall, the repeat unit is a hexagonal hollow ring composed of six interconnected blades. The hollow ring has an inner cavity with diameter of 105 Å. The blade is 180 Å tall. This is substantially shorter than the previous reported value of 235 Å, which was based on a thin-sectioned Spc42p sheets embedded in plastic [7]. The discrepancy of the observed height could be due to reduced resolution in the Z-direction (the end-on direction) in our averaged subtomogram, resulting in a less well-defined boundary for the repeat unit in vertical direction, or alterations caused by the fixation and plastic embedding in the previous study.

**Figure 4.**
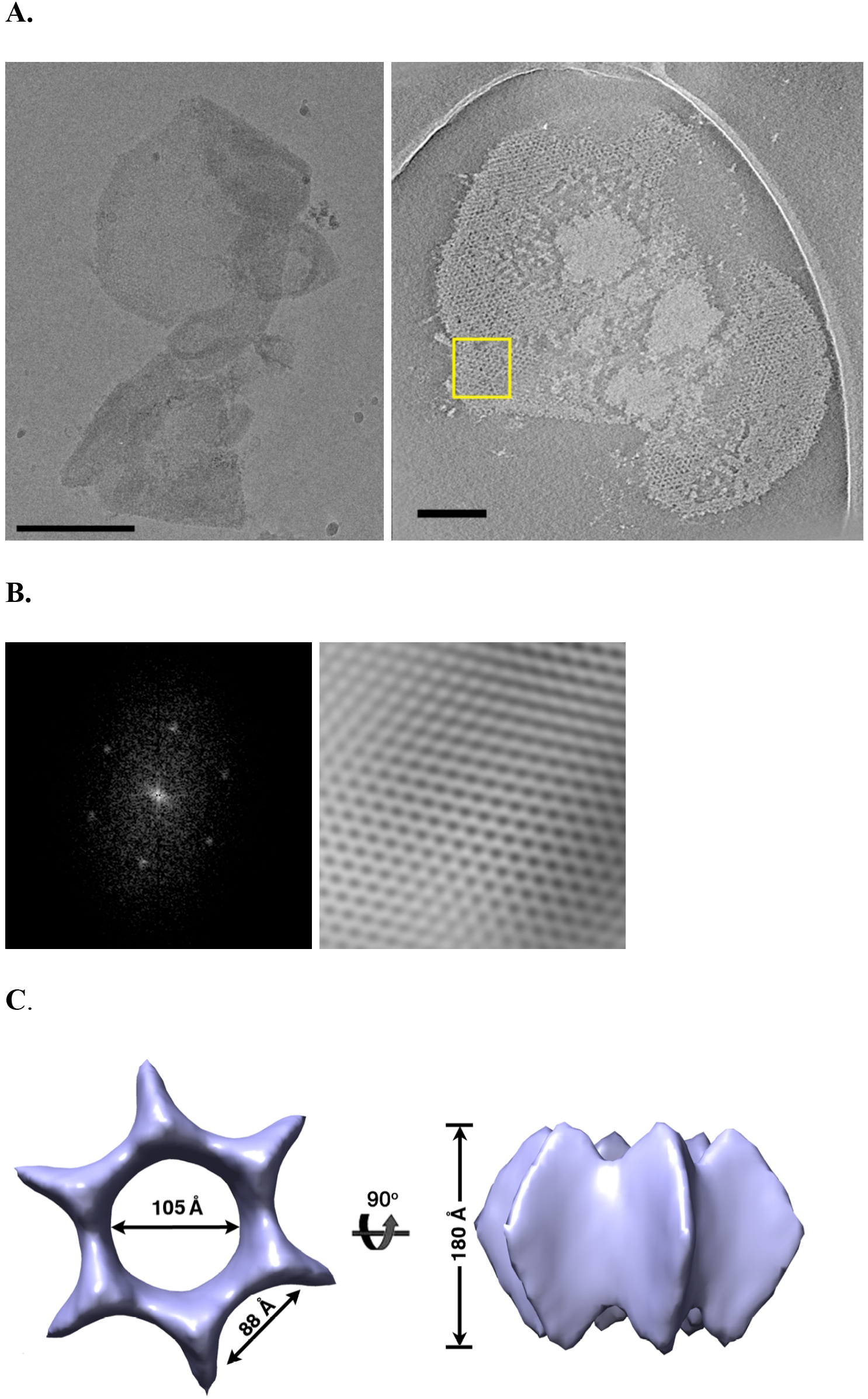
The assembly of sheet by over-expression of Spc42p. (A) A cryo-EM image of an isolated 2D crystal formed by over-expression of Spc42p (Scale, 500 nm) and a central slice from a reconstructed tomogram of the sheet showing ordered assembly. (Scale, 200 nm)
(B) Fourier transform of a boxed area in (A) shows the hexagonal diffraction pattern. The reflections are at spacing of 132 Å. The right side is the filtered Fourier transform. The smearing of the reflections in the diffraction pattern and the Fourier-filtered image indicates local disorder in the crystalline area. Protein (high density) is in white in the right panel.
(C) After subtomogram alignment and averaging, the averaged structure represents the repeat unit in the Spc42p sheet.
(D) A model for the Spc42p assembly based on the averaged structure.
(E) Examples of regular striation pattern in the core of the SPB. The yellow arrow indicates the IL2 layer in each reconstructed SPB where striation is prominent. (Scale,
100 nm)
(F) A reconstructed SPB is shown as tomographic slices at different depth. The yellow arrows indicate the IL2 layer. On the left pattern, the striation spacing in the IL2 layer is ~9 nm, whereas on the right panel, the spacing is ~15 nm. The observation is consistent with the honeycomb model of the Spc42p assembly. (Scale, 100 nm)

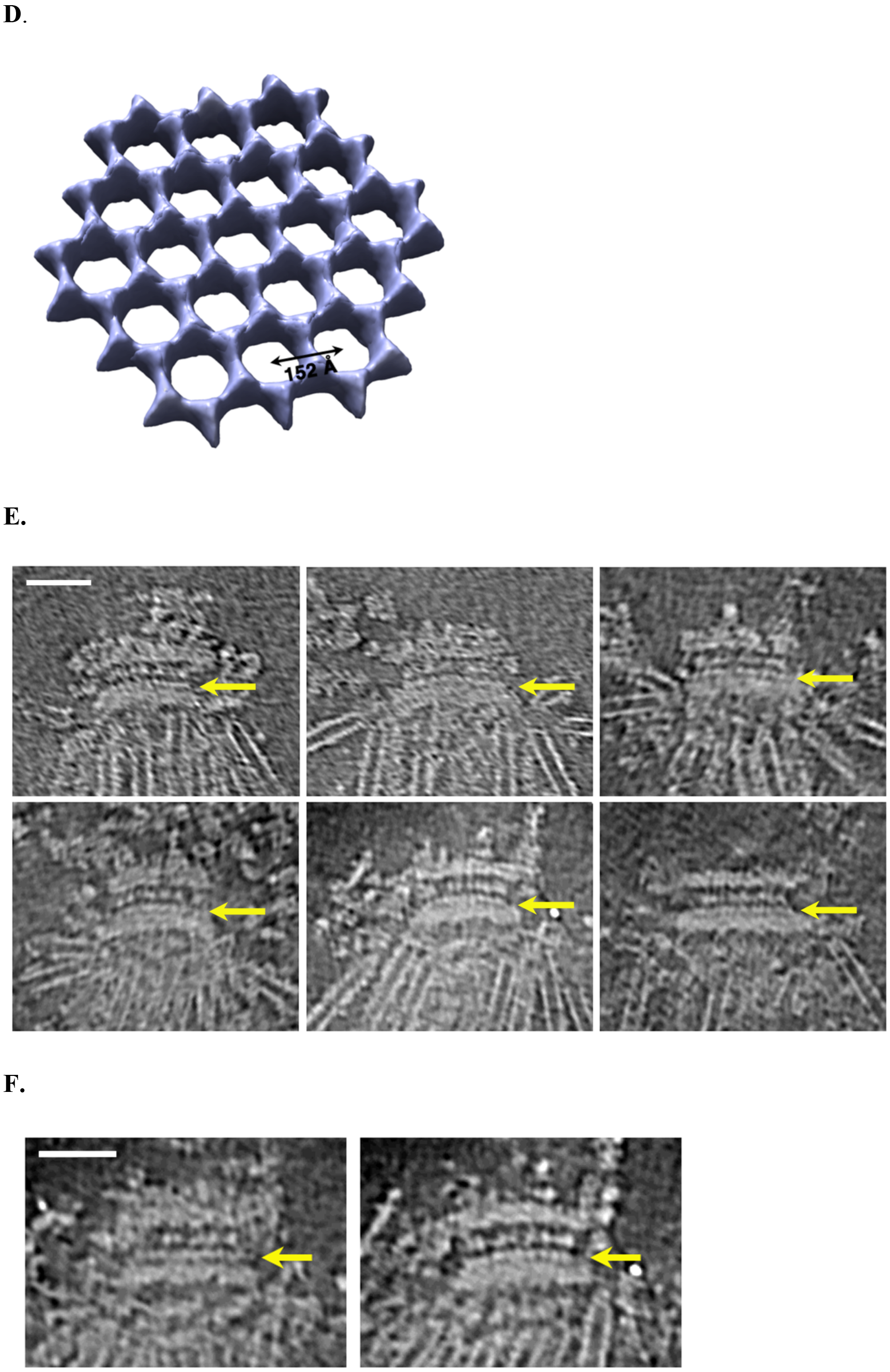

Based on the averaged subtomogram and the position of repeat unit detected in the sheet, we built a model for the Spc42p assembly, as shown in Figure 4D. Overall, the model resembles a honeycomb meshwork. In the model, the distance between centers of a hexagonal repeat unit to its neighbor is 152 Å (152 = 132 / sin60°). Each vertex of hexagon is raised and it is shared by three neighboring hexagons. The wall of hexagon is concaved and shared by two neighbors. Based on the size of hexagon unit, also assuming the average diameter of the SPB core as 160 nm, we estimate there will be ~150 copies of the hexagon unit in each SPB. Furthermore, if we assume 3 copies of Spc42p in each vertex of the hexagon that is shared by three neighboring units, this will bring a total of ~900 copies of Spc42p in a diploid SPB, close to a previous estimation of ~1,000 copies [18], however, a more precise number has to await higher resolution structure of the sheet in the future.

To place this honeycomb structure in the context of the entire SPB assembly, we examined our reconstructed SPB in the tomograms. From 42 volumes examined, 20 showed clear striation in IL2 layers with striation spacing of ~15 nm (Fig. 4E), consistent with the distance between neighboring hexagonal units in the model. Interestingly, depending on the orientation of the SPB and how the tomographic slice cuts through it, the observed spacing in the IL2 layer varies from 9 nm to 15 nm (Fig. 4F). This could be explained by the relative orientation of local hexagon unit in the SPB. In the case of 9 nm, the tomographic slice is likely cutting through the wall of the hexagon, where the spacing between the neighboring ″blades″ along the wall is 88 angstrom (Fig. 4C). The 15 nm spacing observed in the SPB tomogram corresponds to the slice cutting through the center of the hexagonal unit. Together, the regular striation spaces observed in the SPB IL2 layer support the honeycomb meshwork model derived from the overexpressed Spc42p sheet.

The honeycomb structure has high stiffness and strength relative to its weight [39]. The honeycomb-like structure of Spc42p presumably provides SPB the stability and rigidity essential for resisting the pulling and pushing forces during cell division and nuclear migration, and a strong internal scaffold for assembly of other SPB components.

## Conclusion

The purified yeast SPB has provided an excellent opportunity to study the organelle by cryo-ET. In this paper, we have described a comprehensive structural analysis on the yeast SPB, from morphological description of the entire mitotic spindles and the duplicating SPBs to detailed analysis on their local structures and components. This is summarized in Fig. 6.

First we found that the SPB, as a trans-nuclear membrane organelle, is composed of at least eight layers. Based on the measured distances between layers and the regularity within each layer, also by integrating previous published results, we proposed an approximate arrangement of its main components in the SPB. This was followed by a description of three reconstructed tomograms, two mitotic spindles and one duplicating SPB pair. In two mitotic spindles in metaphase, we found the nMTs length spans a large range and was unimodally distributed. There were close interactions between antiparallel MTs. In the duplicating SPB pair, we found the outer layer of the bridge is connected to the CP1 layer of SPB, which is likely to be composed of Spc42p and Spc29p that form the satellite at the beginning of SPB duplication. It also suggests the outer and inner nuclear membrane fuse at the junction between CP1 and CP2 layers. Finally, We used subtomogram averaging to study the assembly of Spc42p. By subtomogram averaging on the reconstructed Spc42p sheet, we provided a 3D map of its repeat unit and built a honeycomb like model for the Spc42p assembly. It is likely to form the main rigid structure of the central plaque and to provide a scaffold for the rest of the SPB assembly. Taken together, the results we present here demonstrate an integration of local structural rigidity and an overall plasticity of functional assemblages in the SPB, which may be common to many other biological systems [40]. Our structural analysis aims to understand the SPB assembly in molecular details. The method of electron cryotomography combined with subtomogram averaging could also be applied to future study of more complex assemblages, such as the mammalian centrosome and other cellular organelles.

## Materials and Methods

### Sample Purification and Preparation

Both SPBs (from *Saccharomyces uvarum*) and Spc42p sheet (from *Saccharomyces cerevisiae*) were purified following previous published procedures [15, 38]. The purified SPB sample, initially in high concentration of sucrose, was first dialyzed at 4°C overnight in a buffer containing 10 mM Bis-Tris/Cl (pH=6.5), 0.1 mM MgCl_2_, 20% (v/v) DMSO. Next day, after mixing with 10 nm colloid gold, the sample was applied onto either a home-made holey carbon grid or a Quantifoil grid (PSI, Inc.) in a humidity chamber, then blotted and plunged into liquid ethane using a home-made plunger or a Vitrobot (FEI, Inc.). Frozen grids were stored in liquid nitrogen before use.

### Electron Microscopy Data Collection

For the purified SPB, single-axis cryo-tomography tilt series were collected on a Tecnai F30 electron microscope (FEI, Inc.) running at 300kV. Images were recorded on a CCD camera (F224, Tietz). Tomographic datasets were collected at a nominal magnification of 15,500, and a defocus value of ~10 μm. The effective pixel size on the images was 9.8 Å. The specimen was tilted from −60° to +60° degree in 2° steps. The accumulated dose for each tilt series was ~80 e^-^/Å^2^.

For the purified Spc42p sheet, single-axis cryo-tomography tilt series were collected on a Tecnai F30 electron microscope (FEI, Inc.) running at 200kV. Images were recorded on a CCD camera (F224HD, Tietz). Tomographic datasets were collected at a nominal magnification of 20,000, and a defocus value of ~7 μm. The effective pixel size on the images was 7.15 Å. The specimen was tilted from −69° to +69° in 1.5° steps. The accumulated dose for each tilt series was ~60 e^-^/Å^2^.

### 3D Reconstruction, Modeling and Subtomogram Averaging

Tomography tilt series were aligned in IMOD [41] by using the 10 nm colloid gold beads as fiducial markers. This was followed by correction for their contrast transfer functions [42]. The 3D reconstructed volumes were calculated by a weighted back-projection program in Priism/IVE [43]. To build models of SPB in mitosis and in duplication, the tomogram volumes were binned 4×4 followed by applying an anisotropic diffusion algorithm to enhance the structural features [44]. Microtubules and the SPBs were manually traced and displayed as surface rendered objects in IMOD [41].

For subtomogram averaging of the Spc42p sheet, an initial model was generated based on a Fourier-filtered image of the 3D reconstructed sheet. In detail, a local patch of sheet was boxed out from a 2D slice extracted from the tomogram volume. Its Fourier transform was calculated, showing the hexagonal pattern. The reflections in the Fourier were manually selected, boxed and their reverse Fourier transformation was calculated by program TRMASK in MRC IMAGE2000 package [45]. This Fourier-filtered image was used to generate an initial 3D model for a hexagonal repeat unit in the sheet. A 3D cross correlation algorithm in Spider [46] was used for searching subtomograms in the 3D volume, followed by subtomogram alignment and averaging. The alignment was carried out iteratively until the structure converged. Care was taken to avoid overlapping region in the sheet before averaging. A total of 660 subtomograms were used for averaging. Since both the diffraction patterns and the Fourier-filtered images strongly suggested hexagonal symmetry, C6 point group symmetry was imposed on the final averaged subtomogram. By splitting the subtomograms into two independent groups, the resolution was estimated as 61 Å at FSC = 0.5 cutoff, though this should be cited with caution since the averaged structure is substantially anisotropic because of dominance of end-on views of the sheet in ice.

**Figure 5.**
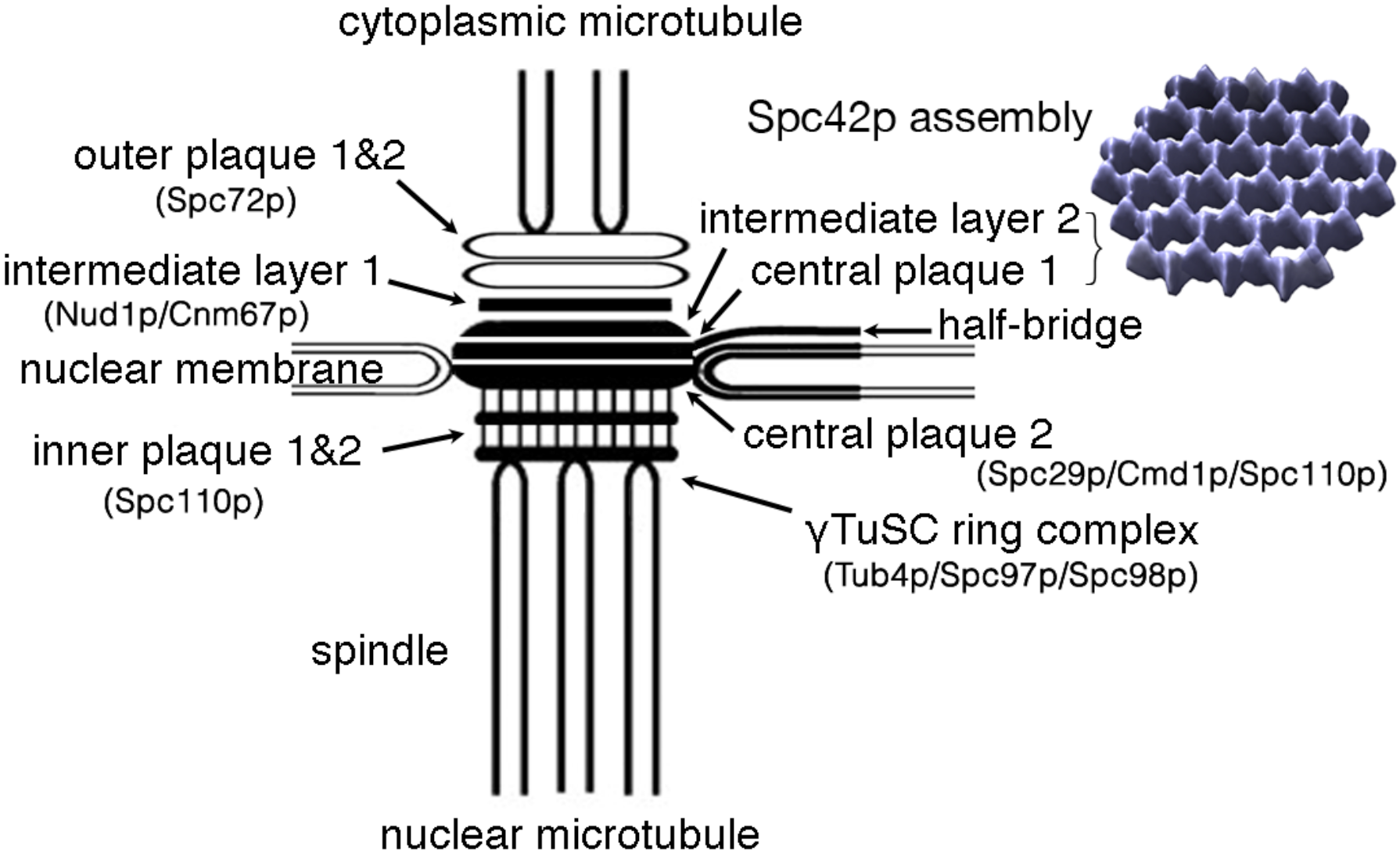
A schematic diagram to summarize the current study of the SPB. Eight layers of the SPB are labeled. The averaged structure of spc42p layer that forms the SPB core, are displayed as surface rendered image. For display purpose, the distances between layers and the sizes of averaged structures are not in proportion.

## Acknowledgements

The authors thank J. V. Kilmartin for providing the enriched SPB sample, R.A. Crowther, R. Henderson for their advice and encouragement during the course of this study conducted at MRC-LMB. We thank J. Berriman and S.X. Chen for help on electron microscopy and imaging. SL was recipient of a HFSP long-term fellowship and a MRC career development fellowship. This work was supported by MRC.

## Supplemental Material

### 1. Supplemental Figures

**Figure S1.**
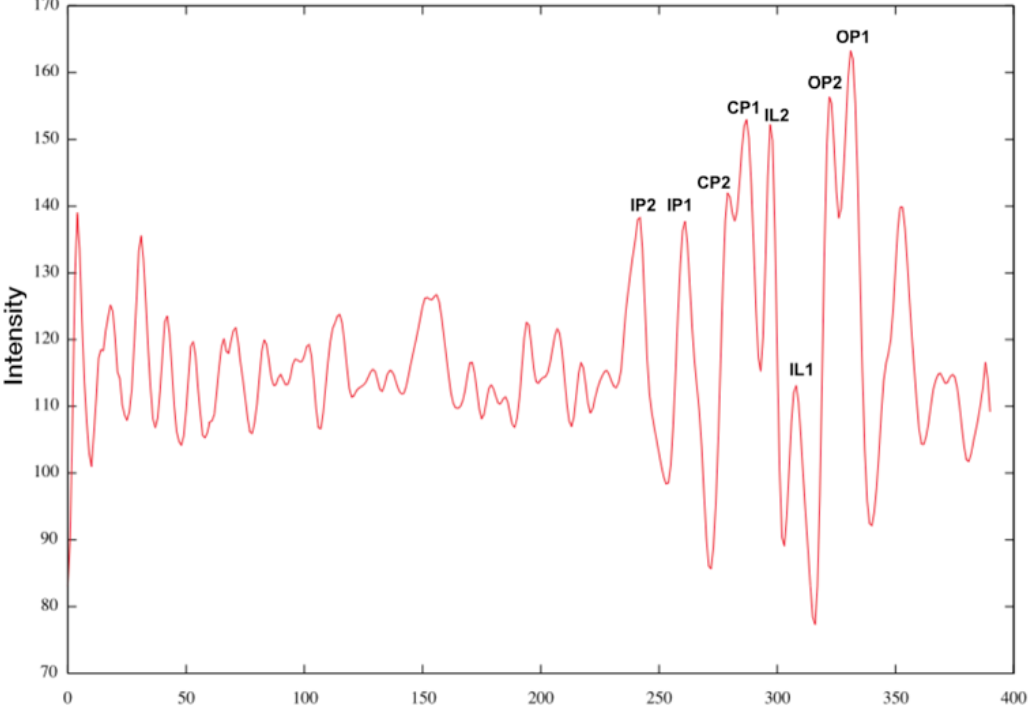
Histogram of radial averaged intensity based on tomographic reconstructed SPB (Horizontal axis is in the unit of pixel. 19.6 Å/pixel).

### 2. Supplemental videos

**Video 1**. An aligned tomographic tilt series with a SPB embedded in vitrified ice.

**Video 2**. A reconstructed 3D volume containing a SPB and a bundle of nMTs.

**Video 3**. A reconstructed 3D volume of a yeast spindle in early mitosis.

**Video 4**. A second mitotic spindle reconstructed by tomography

**Video 5**. Tomographic reconstruction of a pair of SPBs in duplication. Two SPBs are linked by a bridge structure.

